# Peptide-centric proteomics reveals molecular subtypes of sporadic Alzheimer’s disease

**DOI:** 10.64898/2026.11.12.766321

**Authors:** Deanna Plubell, Gennifer Merrihew, Jea Park, Karynne Patterson, Joshua D. Smith, Chris Hsu, Christine C Wu, C. Dirk Keene, Thomas Montine, Michael J MacCoss

## Abstract

Alzheimer’s disease (AD) is a complex, age-related disease that continues to increase in incidence worldwide. Although some cases are caused by known pathogenic mutations, the majority of cases are sporadic, with uncertain etiology and commonly complicated by comorbid diseases. To better understand the molecular landscape of AD we constructed a unique brain sample set free from neuropathologic comorbidities. This cohort consists of autosomal dominant AD dementia, sporadic AD dementia, cognitively normal individuals with high AD neuropathologic change - a condition called resilient to AD, and cognitively normal cases with no or minimal AD neuropathologic change. Through quantitative mass spectrometry proteomics we obtain measures of peptides mapping to Aβ and to tau, capturing differences in the histopathologic burden of disease. Correlations of these peptide abundance differences within sporadic AD dementia cases in the superior and medial temporal gyrus indicate the presence of two separate sporadic AD phenotypes. Additional differences are characterized at a peptide level between sporadic AD dementia and autosomal dominant AD dementia in the inferior parietal lobe and caudate nucleus. These findings indicate both shared and distinct molecular features in AD influenced by cause and age of onset of disease.

## INTRODUCTION

Alzheimer’s disease (AD) is a major global public health threat. In contrast to ischemic heart disease, stroke and several forms of cancer, AD dementia is increasing as a cause of death, of years lived with disability, and of disability-adjusted life years^1^. AD is a chronic illness whose clinical expression typically begins with memory impairment and culminates as dementia following years, if not decades, of brain regional neuronal injury, response to injury, consumption of reserve, and exhaustion of compensation. AD is fundamentally characterized neuropathologically by the stereotypical accumulation of amyloid-β-containing plaques and pathologic tau-containing neurofibrillary tangles. Since animal models of AD still struggle to capture the complexity of the human condition, determining the molecular profile specific to AD in its various forms in people independent of common comorbidities will be essential to efforts to develop tailored therapies that specifically target the molecular mechanism(s) of AD.

A small proportion of people with AD have a known genetic cause that usually is inherited as an autosomal dominant (ADAD) trait with high penetrance, although de novo variants have also been described^2–4^. Almost all known genetic causes of AD occur in the genes encoding presenilin-1 (PSEN1), presenilin-2 (PSEN2), or amyloid precursor protein (APP)^5–7^. Pathogenic variants in these genes result in the aberrant processing of APP to amyloid-β isoforms, likely due to a change in binding site affinity or specificity, and are proposed to trigger regionally stereotypical neurofibrillary degeneration through a number of potential mechanisms^3,4^. Importantly, accumulation of amyloid-β peptides and pathologic tau also occurs in AD with no known genetic cause, called sporadic AD^5,8^. Despite the shared neuropathologic hallmarks, there are often differences in age of onset, rate of progression, and involvement of comorbidities between ADAD and sporadic AD (SAD)^8–10^. By far, the strongest genetic risk for SAD is the ε4 allele of the apolipoprotein E gene (*APOE*)^11,12^.

Existence of forms of AD with known genetic causes or risk, without known genetic underpinnings, and with pathologic comorbidities, highlight the potential for multiple molecular drivers and perhaps multiple pathogenic pathways involved in the most common form(s) of cognitive impairment and dementia. To better understand the molecular signatures of AD we constructed a unique human brain sample set with ADAD dementia, SAD dementia, high cognitive function with high AD neuropathologic change (resilient to AD or RAD), and high cognitive function with no/minimal AD neuropathologic change (healthy control or HC); importantly, all cases were selected pathologically to exclude comorbidities so that our data reflect only the contributions of AD. Using data-independent acquisition mass spectrometry proteomics, we examined differences between cohort conditions at a peptide and protein level in four brain regions^13^.

## SUMMARY OF MAJOR FINDINGS

Peptide measurements significantly correlated with known pathological markers of disease and separated SAD into two subgroups based on the superior & middle temporal gyrus proteome. These subgroups are distinct primarily in their age of death, indicating a difference in the severity of disease. Additional differences between ADAD and SAD are described in the inferior parietal lobe and caudate nucleus proteomes, and comparisons across brain regions demonstrate similar signatures of disease in both cerebral cortical regions. This study highlights the use of characterizing the proteome of AD to further understand clinically meaningful subgroups.

## RESULTS

### Unique and shared proteome signatures across brain regions

Protein groups with significantly altered abundances between the four diagnostic groups were compared across the four brain regions that are variably impacted by the two hallmark features of AD. The inferior parietal lobule (IPL) has the most significantly different proteins (2891 total), followed by the caudate nucleus (CN) (2566 total), superior and medial temporal gyri (SMTG) (2558 total), and hippocampus (144 total). In each analysis, the comparison between ADAD and both HC and RAD had the largest number of significant differences (Supplemental table). The largest overlap in significant proteins is between the two isocortical regions (IPL and SMTG), with 842 shared proteins (Figure 1a). Beyond those shared proteins, 1072 are uniquely altered in CN, 753 proteins are uniquely altered in IPL, and 546 are uniquely altered in SMTG. There are an additional 830 proteins significantly altered among IPL, SMTG, and caudate, and only 88 proteins significantly altered among all four brain regions. Proteins uniquely different in each brain region are enriched for different gene ontology terms. Differential proteins unique to the caudate are enriched in terms primarily related to cytoplasmic translation, intracellular and vesicle-mediated transport, endocytosis, and membrane organization (Figure 1b). Proteins unique to SMTG are enriched in mitochondrial gene expression, muscle contraction, and myelination (Figure 1c). For the IPL, unique differential proteins are enriched in terms primarily related to RNA splicing and gene expression, lysosomal, Golgi vesicle and endosomal transport, and regulation of actin filaments (Figure 1d). The few ADAD hippocampus samples resulted in no significant enrichment for gene ontology terms.

**Figure 1.**
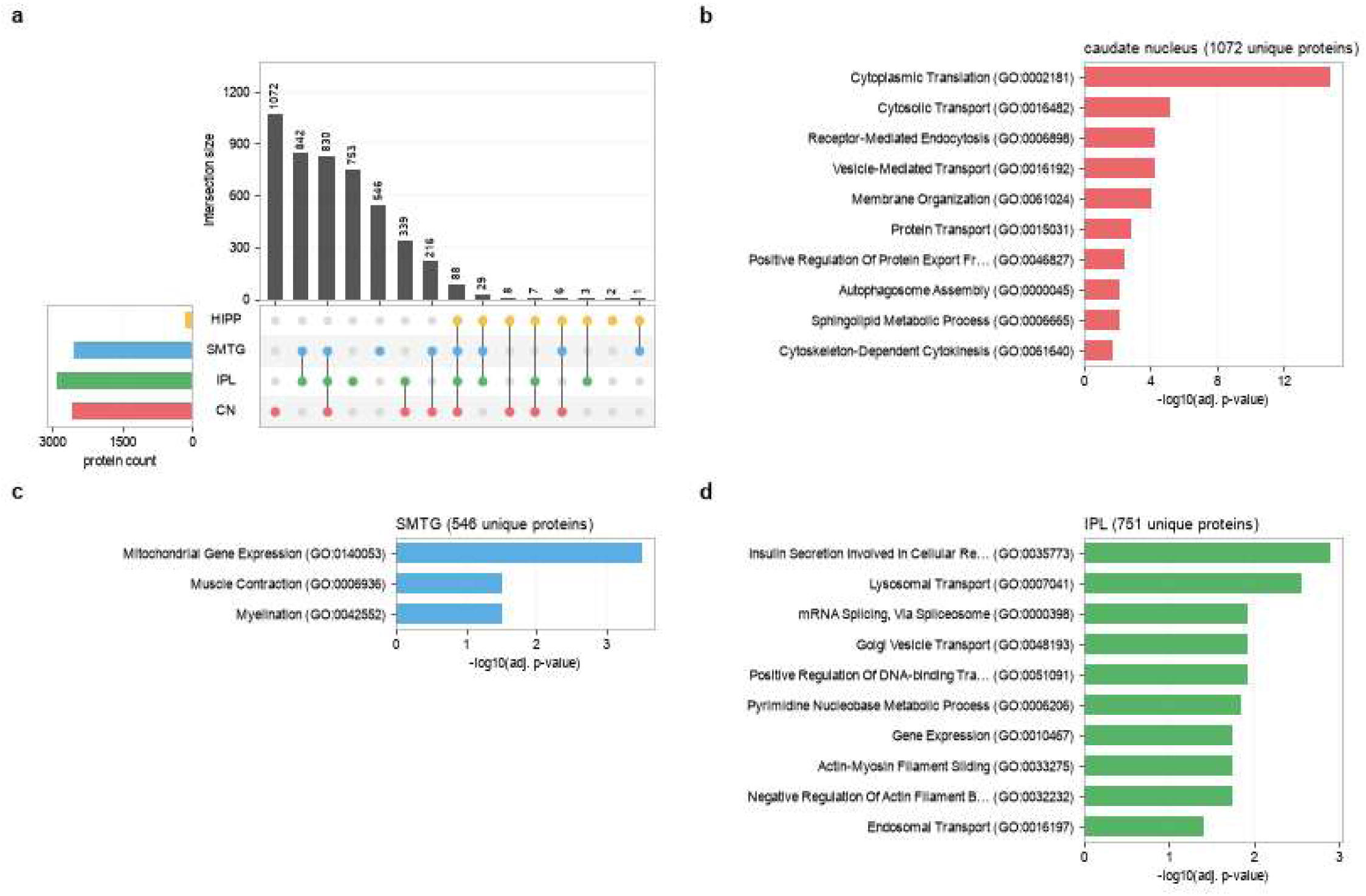
Differential protein abundance profiles among brain regions. A) The overlap in proteins with significant differences across diagnostic groups for each brain region. Gene ontology terms enriched in the proteins with significantly altered abundance in B) the caudate nucleus proteome, C) the SMTG proteome, and D) the IPL proteome.

### Proteomics further differentiates within neuropathologic staging

Two tryptic peptides with sequence mapping to the amyloid-β region of Amyloid precursor protein (APP) are quantified. These peptides have very high relative intensity compared to the other APP peptides, and the expected differences across groups with the HC having the lowest abundance followed by the RAD, SAD, and ADAD. The amyloid-β tryptic peptide LVFFAEDVGSNK (APP residues 688-699; amyloid-β residues 17-28) in all brain regions shows more separation between conditions with the same histopathologic assessment for tissue density of neuritic amyloid-β plaques (CERAD score as C0 to C3), compared to CERAD scores in the same condition (Figure 2a). The distinction between conditions is most apparent in individuals scored with the highest tissue density of plaques (C3), with ADAD having higher abundances compared to the SAD or both RAD and HC. Similar trends are also observed for tau with abundances across regions reflecting the higher neuropathologic burden in ADAD. Interestingly, the peptide abundance of IGSLDNITHVPGGGNK (tau residues 354-369) better distinguishes between SAD and HC and RAD in the hippocampus, SMTG, and IPL. Like the amyloid-β tryptic peptide, the tau peptide IGSLDNITHVPGGGNK across brain regions shows a slight separation between diagnosis groups within histopathologic staging of neurofibrillary degeneration (NIA-AA B score, B0 to B3). These differences are observed primarily between SAD and ADAD in cases with a high B score (B3).

**Figure 2.**
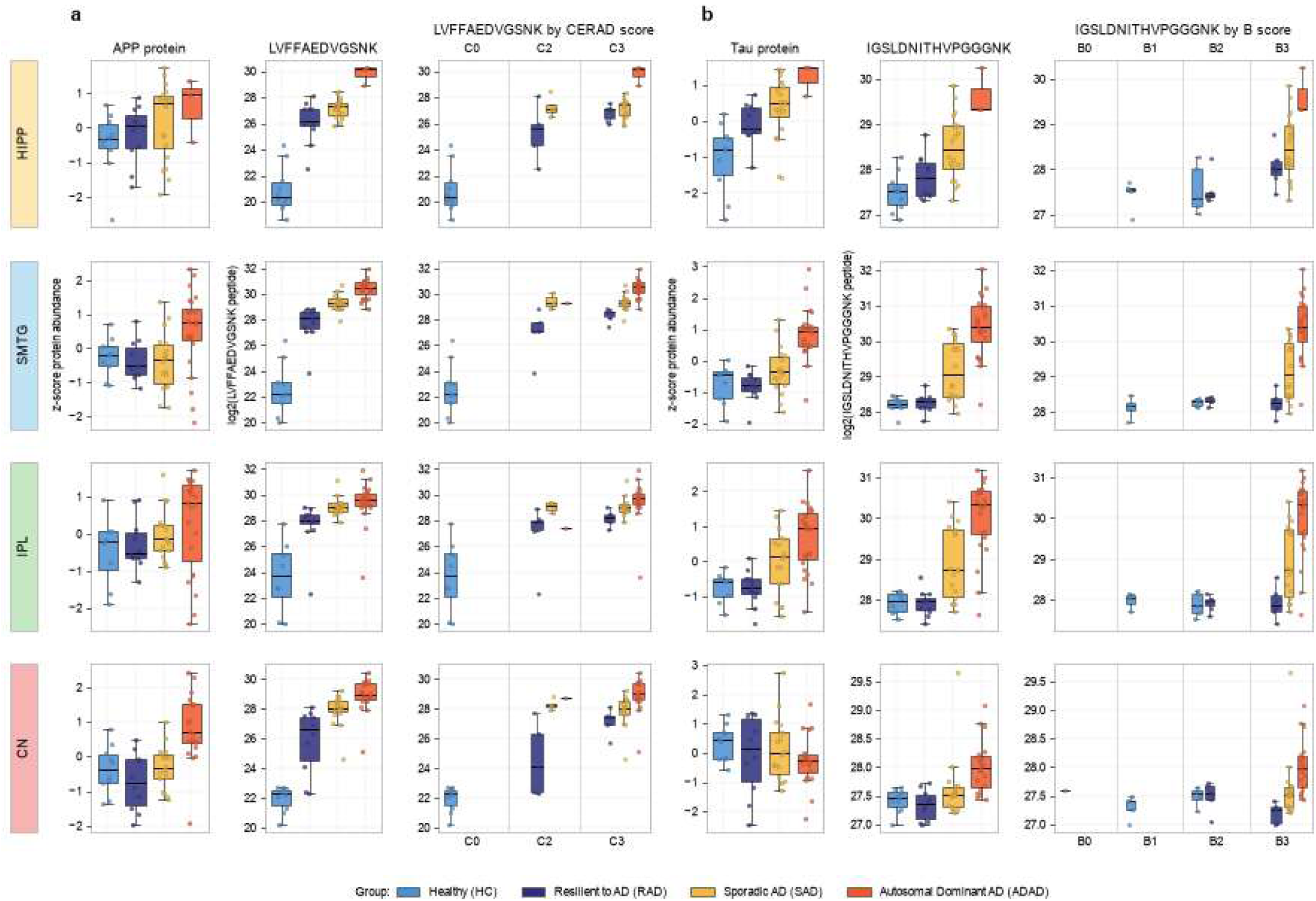
Amyloid precursor protein (APP) and tau peptide measurements distinguish between diagnosis groups. A) Calculated APP protein abundance z-score normalized for comparison to peptide abundances, and the log2 peptide abundance for the peptide LVFFAEDVGSNK mapping to residues 688-699 of APP (residues 17-28 of Aβ), also stratified by individuals CERAD score for neuritic plaques. B) Calculated tau protein abundance z-score normalized for comparison to peptide abundances, and the log2 peptide abundance for the peptide IGSLDNITHVPGGGNK mapping to residues 354-369 of tau, also stratified by NIA- AA B score for neurofibrillary degeneration. Condition is indicated by bar color with healthy control (HC) far left, then resilient to AD (RAD), sporadic AD dementia (SAD), and ADAD.

### Peptides co-varying with amyloid in SMTG

The Aβ peptide LVFFAEDVGSNK abundance varied by neuropathologic changes across all brain regions and was selected as a brain region specific continuous proxy of AD neuropathologic change to identify other tryptic peptides with associated abundances (Figure 3). In SMTG samples, 1626 peptides mapping to 601 proteins were significantly correlated with the LVFFAEDVGSNK abundance based on a Spearman’s rank correlation with a Bonferroni corrected p-value < 0.05. Of the correlated peptides 51.7% (840) were positively correlated with Aβ 17-28 abundance and 48.3% (786) were negatively correlated. Of the proteins represented in the positively correlated peptides, the gene ontology enrichment analysis indicated an enrichment in terms related to immune cell migration and adhesion, integrin signaling, actin cytoskeleton reorganization, proteasomal protein degradation, and phagocytosis. Proteins represented in the negatively correlated peptides are enriched in terms primarily related to aerobic respiration and pyruvate metabolism, vesicle-mediated transport, and neuron projection morphogenesis.

**Figure 3.**
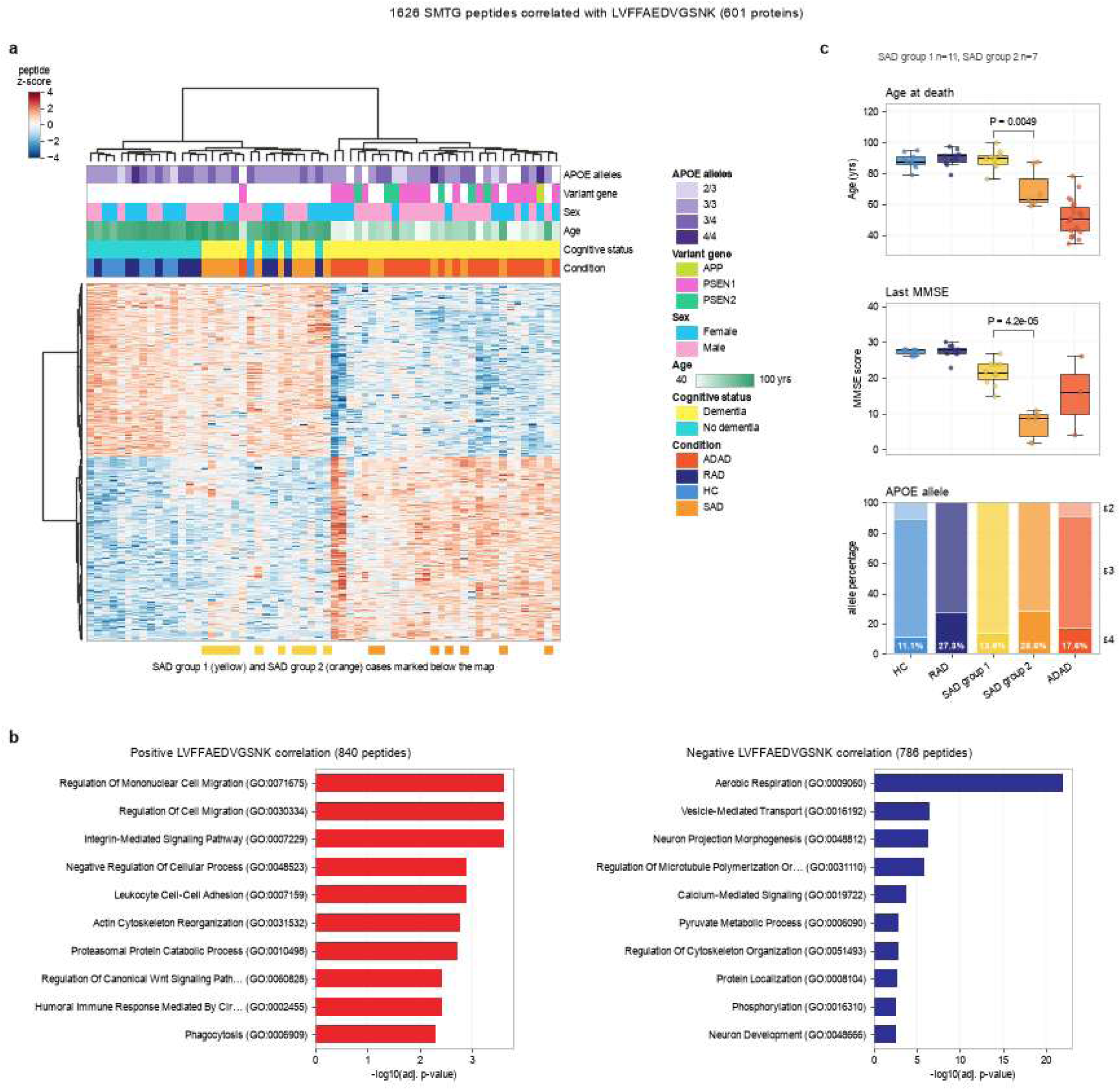
Peptides correlated with APP LVFFAEDVGSNK in SMTG distinguish between subgroups of sporadic AD (SAD). A) Heatmap of significantly correlated peptide abundances (adj. P-value <0.05) with hierarchical clustering of both cases and peptides. B) Gene ontology enrichment analysis of positively and negatively correlated peptides. C) Characteristics for diagnostic groups with SAD separated into subgroups (n=11 SAD group 1, n=7 SAD group 2,) with significant differences between the SAD subgroups for age (p=0.0049) and last MMSE (p=4.2e-05). Condition is indicated by bar color with HC far left, then RAD, SAD group 1, SAD group 2, and ADAD.

For the peptides significantly correlated with Aβ peptide LVFFAEDVGSNK in SMTG, hierarchical clustering identified two groups with one composed of all but one of the ADAD samples and the other composed of the HC and RAD samples and the remaining ADAD sample. Interestingly, the 18 SAD cases separated across these two clusters, with 11 SAD subclustering with the HC and RAD samples (SAD group 1) and 7 SAD subclustering with the majority of the ADAD (SAD group 2) samples. Since our analysis clearly separated the SAD samples into two distinct subclusters, we investigated the features underlying this separation. Any artifacts of the batches used for sample preparation and acquisition were ruled out. From the clinical characteristics we observed that the individuals in SAD group 2 were significantly younger at the time of death compared to SAD group 1 individuals (p=0.0049, Figure 3) but still with some overlap in age ranges. Although there are many possible factors that influence the interval between acquiring the last MMSE and autopsy, one is the challenges of highly impaired individuals being able to accommodate a clinical research evaluation. With these limitations in mind, the last MMSE affiliated with each sample is also significantly different between the two sporadic groups (p=4.2e-05); with complete separation between those that cluster with ADAD and those that cluster with the HCF. Since the post-mortem interval tended to be longer for most of the ADAD cases, the post-mortem interval between the two SAD clusters was tested, but found to be not significantly different (p=0.40, Supplemental table 1). *APOE* genotype of the SAD samples show an increased percentage of ε4 alleles in the ADAD-like SAD group 2 samples, but the cohort size limits meaningful interpretation of allele frequency.

Based on the differences in proteome and age in the two SAD sample clusters we investigated whether there was a genetic misclassification of these samples. Exome sequencing was performed on all 18 SAD samples that had their SMTG tissue analyzed by proteomics. Sequence alignment and variant calling was performed for genes known to cause ADAD. For the genes analyzed, a total of 8 variants were called in 7 individuals, in ABCA7, SORL1, and APP (Table 1). No variants were found in either PSEN1 or PSEN2. Most of the variants found were missense, while one was a synonymous variant and one resulted in a frameshift (p.Leu1403ArgfsTer7) in ABCA7.

**Table 1.** Variants called in genes implicated in AD. Sporadic AD cases were exome sequenced, with variants called for *APP, PSEN1, PSEN2, ABI3, ABCA7, SORL1*, and *TREM2*. *Indicates found in the same individual.

| Group | Gene | Type | Allele count (gnomAD) | Allele frequency (gnomAD) | Clinical significance (ClinVar) | Age (yrs) | Sex | APOE genotype | StudyName |
|---|---|---|---|---|---|---|---|---|---|
| **SAD group 2** | *ABCA7* | frameshift | 234 | 8.80E-04 | Conflicting | 88 | Male | ε3/ε3 | UW ADRC |
|  | *SORL1* | missense | 7 | 2.78E-05 | NA | 61 | Male | ε4/ε4 | UW ADRC |
| **SAD group 1** | \**ABCA7* | missense | 55 | 1.96E-04 | NA | 77 | Male | ε3/ε3 | ACT |
|  | \**ABCA7* | missense | 55 | 1.99E-04 | Uncertain significance | 77 | Male | ε3/ε3 | ACT |
|  | *ABCA7* | synonymous | 1 | 3.59E-05 | NA | 86 | Male | ε3/ε4 | UW ADRC |
|  | *SORL1* | missense | 715 | 2.54E-03 | Likely benign | 91 | Female | ε3/ε4 | UW ADRC |
|  | *SORL1* | missense | 715 | 2.54E-03 | Likely benign | 100 | Male | ε3/ε3 | ACT |
|  | *APP* | missense | 28 | 1.11E-04 | Uncertain significance | 89 | Female | ε3/ε3 | ACT |

*\*Same individual*

### Proteomics signatures separate SAD groups across regions

The same separation of the SAD samples in the SMTG also can be observed in proteome-wide analyses for the other brain regions (Figure 4). Principal component analysis on the peptides correlated with Aβ peptide LVFFAEDVGSNK (Hipp n=34, SMTG n=1626, IPL n=291, Caudate n=71) cluster the SAD group 2 as expected, but principal component analysis on all measured peptides also separates the dominant-like SAD group 2 from dominant-unlike SAD group 1 (Supplemental Figure 3). Neither principal component analyses clearly separated SAD group 1 from the high cognitive function individuals (HC and RAD). For Aβ and tau tryptic peptides, the Aβ peptide LVFFAEDVGSNK is relatively similar between the SAD group 1 and group 2, while both tau SGYSSPGSPGTPGSR (residues 195-209) and tau IGSLDNITHVPGGGNK (residues 354-369) have different distributions between the two SAD subgroups. There are still differences between SAD group 1 and group 2 in tau and ApoE peptides in IPL, but the Aβ LVFFAEDVGSNK correlated peptides in IPL do distinctly separate SAD cases in the same manner as in the SMTG (Figure 4). Signatures of dominant-like and dominant-unlike SAD are not detected in the caudate nucleus samples, either in the correlated peptides (Figure 4a) or in differential abundance analysis. Peptides in Aβ, tau, and ApoE also do not have altered abundance between the SAD subgroups. Differential abundance testing was also performed with the two SAD subgroups and the ADAD, RAD and HC. These results recapitulate the clustering analysis, with the largest number of differential peptides in SMTG between ADAD and SAD group 1 (14,694), followed by SAD group 2 and SAD group 1 (11,805), ADAD and HC (11,629), and ADAD and RAD (10,464), while only one peptide differed between ADAD and SAD group 2.

**Figure 4.**
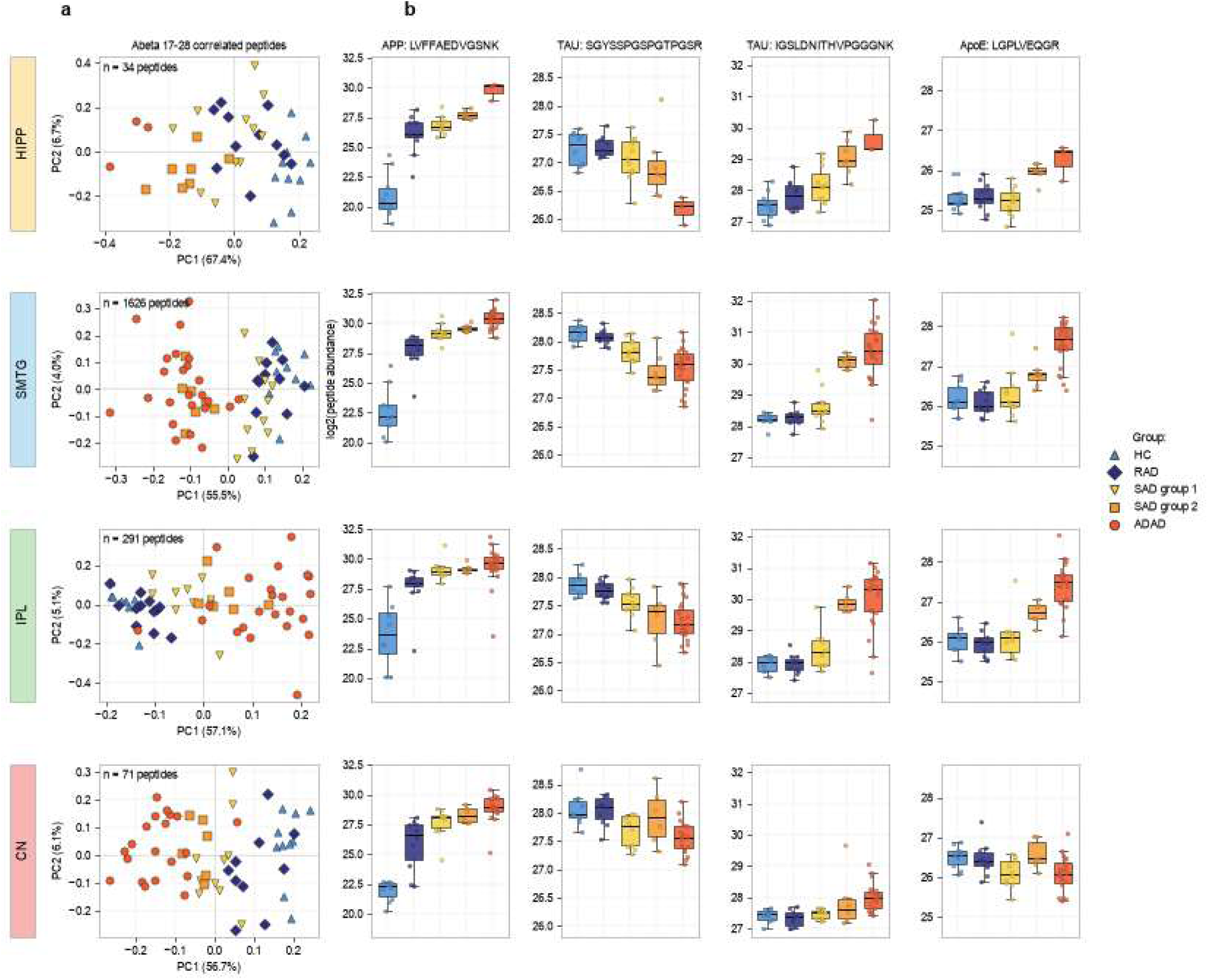
SAD subgroups are most distinct in SMTG. A) Principal component analysis of APP LVFFAEDVGSNK (residues 17-28 of Aβ) significantly correlated peptides (adj. P-value <0.05) for each brain region. B) Condition is indicated by bar color with HC far left, then RAD, SAD group 1, SAD group 2, and ADAD. Tau peptide SGYSSPGSPGTPGSR corresponds to residues 195-209 and IGSLDNITHVPGGGNK corresponds to residues 354-369, while ApoE peptide LGPLVEQGR corresponds to residues 199-207.

### Amyloid precursor protein is increased in abundance in ADAD across several brain regions

We quantify 26 unique peptides that map to APP, with 21 detected in all four brain regions (Figure 5a, Supplemental Figure 4). Peptides span the APP sequence, including two peptides mapping to the Aβ peptide region, spanning the gamma secretase site (HDFS, LVFFAEDVGSNK). ADAD cases overall have higher APP peptide abundance in hippocampus, SMTG, and caudate regions (Figure 5). This is also observed when all peptides are rolled up to a single protein abundance (Figure 2a), but is not only due to the majority of the signal coming from Aβ peptide LVFFAEDVGSNK. APP peptide abundance further separates both the SAD groups, with individuals in SAD group 1 tending to more closely reflect the HC and RAD abundances in the hippocampus and SMTG, but not the IPL or caudate. ADAD individuals with a PSEN1 variant also generally have higher APP peptide abundance than those with a PSEN2 variant, while not statistically significant (Supplemental Figure 1). There does not appear to be a consistent trend in either APP or the LVFFAEDVGSNK peptide abundance by APOE allele within each condition (Supplemental Figure 2).

**Figure 5.**
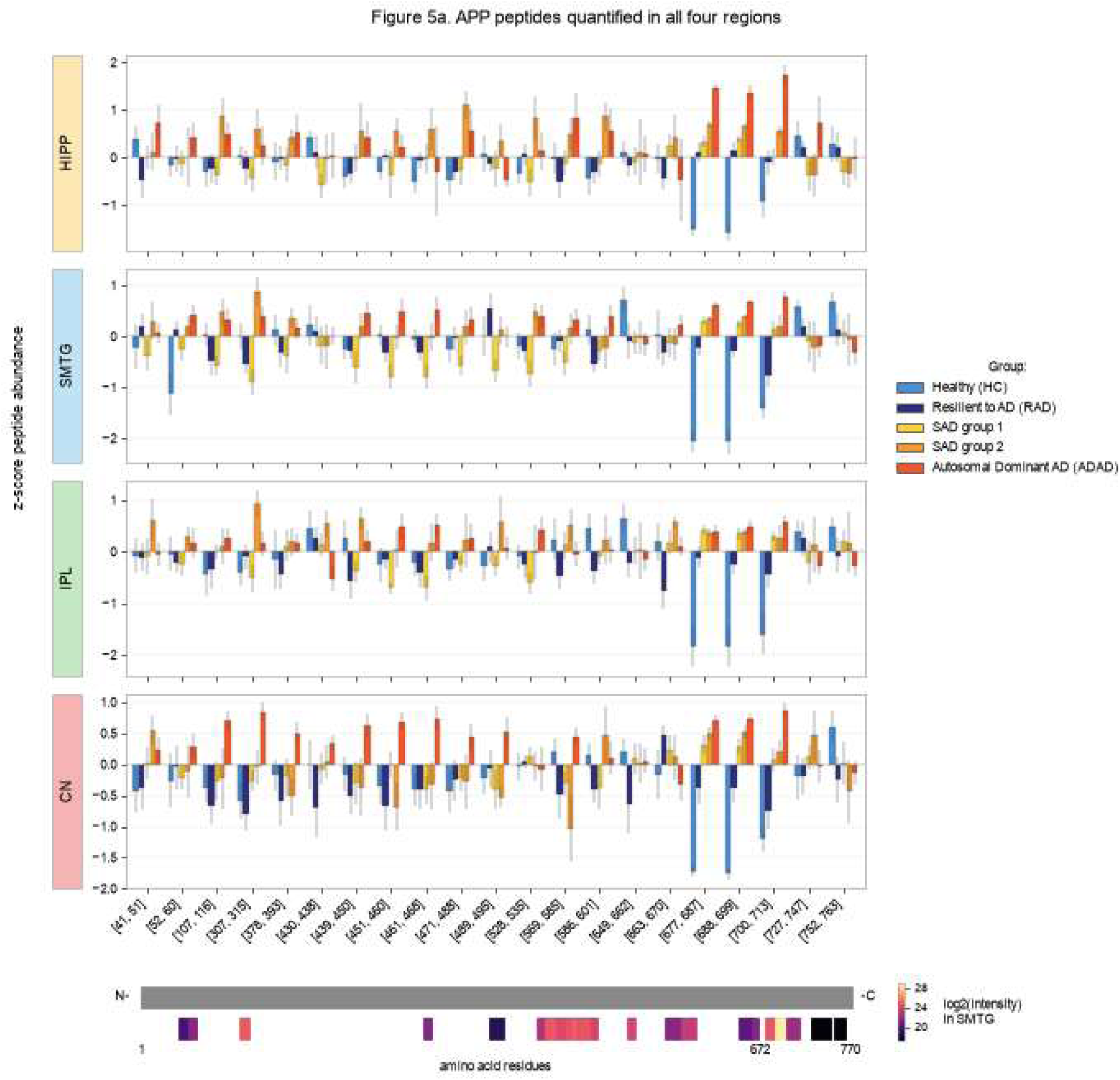
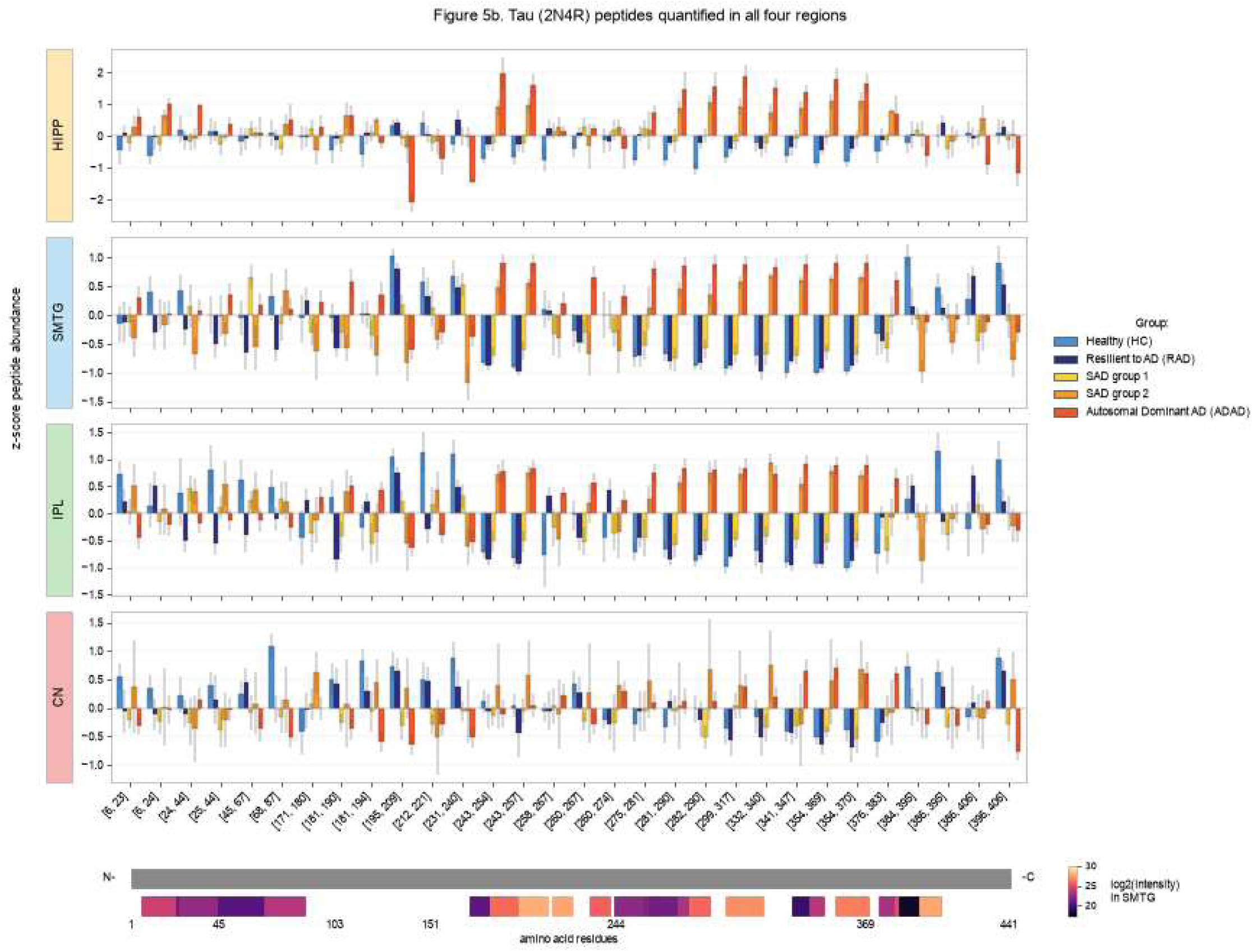
Comparison of APP and tau peptide abundances across brain regions. Tryptic peptides quantified in all four brain regions mapping to A) APP and B) tau (2N4R isoform, P10636-8) are z-score normalized across samples and plotted by mean +/- se per condition from N- to C-terminus. Condition is indicated by bar color with HC far left, then RAD, SAD group 1, SAD group 2, and ADAD. Protein is represented from N- to C-terminus, with peptides mapped and colored by their mean log2(signal intensity) in SMTG.

### Tau hyperphosphorylated region and microtubule binding domain have inverse profiles across samples

We quantify 48 unique peptides mapping to microtubule associated protein tau (Supplemental Figure 5). Of these, 35 map to the 2N4R isoform, and 30 of those were quantified in all four brain regions (Figure 5b). The quantified peptides span most of the 2N4R tau sequence isoform (441 aa, UniprotKB P10636-8); with peptides mapping to all variable domains. Abundance profiles are most similar between the SMTG and IPL, with some differences in the hippocampus and caudate (Figure 5b). Most peptides mapping to the microtubule binding regions in R2-R4 (residues 275-368) have similar abundance profiles; HC has the lowest abundance, followed by RAD, SAD group 1, SAD group 2, and ADAD. The peptide mapping to R1 (260-267) has the same abundance trend in the hippocampus and to a lesser extent in the SMTG, but does not share the same abundance trend across conditions in the IPL or caudate. Peptides mapping to the proline-rich regions of P1 and P2 in the SMTG and IPL have the inverse abundance trends to the R2-R4 peptides, with the lowest abundances in ADAD, followed by SAD group 2, SAD group 1, RAD, and HC. The SAD subgroups are separated by peptides in both tau regions primarily in the SMTG and IPL, but not as distinctly in hippocampus, and no apparent separation in the caudate nucleus. Two c-terminus tau peptides (386-395 and 396-406) also demonstrate abundance profiles more similar to the P1 and P2 regions, particularly in the SMTG and IPL regions. Thirteen additional peptides are detected and quantified in several of the brain regions that do not map to the 2N4R isoform; instead mapping to the isoform referred to as PNS-tau (Supplemental Figure 5). These were found due to the inclusion of the PNS-tau 758 aa sequence in the human proteome FASTA as the canonical MAPT sequence (UniprotKB P10636-1).

## DISCUSSION

We find distinct differences at a proteome level among four different diagnostic groups and among four different brain regions. The four diagnostic groups include HC, RAD, SAD, and ADAD – all carefully evaluated to exclude common comorbidities thereby enabling us to focus on the proteomic signature of AD without confounding. We deliberately selected four regions of the brain with varying cell composition, tissue structure, and neural circuits subserved as well as varyingly impacted by AD neuropathologic change, to gain deeper insights into proteomic signatures. Finally, we employed cutting-edge proteomic techniques, multiple robust technical controls, and sophisticated analyses to generate high-quality proteomic data unencumbered by challenges recently highlighted for other proteomic approaches. Our focus on quality of case annotation and data generation necessarily limits the number of samples that meet our inclusion/exclusion criteria and rigorous analysis. We recently described this approach using a subset of these samples and protein level data to compare protein signatures of RAD versus HC and SAD^14^. Here we focused on peptide level changes and critically included ADAD samples.

Increased regional tissue abundance of Aβ is a hallmark of AD. The Aβ sequence mapping peptides are increased as expected based on neuropathologic selection criteria for this sample set. Unsurprisingly, despite ordinal histopathologic ranking of Aβ plaque density being the same across many AD cases, the abundance of Aβ peptides distinguished among the diagnostic groups, confirming at the peptide level our protein level analysis of RAD versus HC and SAD, and importantly extending these data to include ADAD. This is likely due to the more precise and comprehensive measures by mass spectrometry than by immunohistochemistry, allowing us to expand the dynamic range and improve sensitivity, especially in cases with highly abundant Aβ as occurs in ADAD. Moreover, while neuropathologic assessments evaluate higher order aggregates like Aβ fibrils, proteomic measurement of tryptic peptides mapping to Aβ encompass all forms of Aβ and the Aβ sequence from intact APP. However, it is likely that the majority of the Aβ tryptic peptides signal is coming from Aβ peptides due to the observed difference in relative intensity of these peptides compared to other APP peptides.

Unlike the expected differences in Aβ, the increased abundance of APP in ADAD is not well described. We detect peptides across the whole APP sequence to varying degrees in all brain regions, and find most peptides are increased in ADAD compared to SAD in the hippocampus, SMTG, and caudate nucleus. Previous work has demonstrated that the entire APP protein is increased in some causal variant carriers, specifically with *APP* causal variants^15,16^; however, we observed an increase that is relatively consistent across all causal variants. The mechanisms that increase APP abundance in ADAD are not clear, with a preliminary trend of APP being slightly further elevated in *PSEN1* variant cases compared to *PSEN2* carriers. Similar to APP, ApoE is also distinctly increased in ADAD in hippocampus, SMTG, and IPL, but not in the caudate, perhaps related to ApoE binding to aggregated forms of Aβ. There are some preliminary trends in Aβ abundance related to *APOE* genotype; however, our sample size is too small to powerfully address this intriguing observation.

An unexpected finding was the distinct differences in SMTG proteomes within the SAD cases. These differences were primarily observed in the peptide abundances that significantly correlated with Aβ 17-28 abundance, but were also observed by clustering performed on the whole proteome. Because we had a subgroup of SAD cases that had proteomic profiles more similar to ADAD we determined if those individuals carried causal variants in *APP, PSEN1*, or *PSEN2* that had not been considered previously. Additionally, several genes have been identified as implicated in AD through GWAS; specifically, *TREM2, ABCA7, SORL1,* and *ABI3*^17–23^. These 7 genes were analyzed by whole exome sequencing for the 18 SAD cases. While some variants were found in ABCA7, SORL1, and APP, none had known pathogenic status. One potentially functionally significant variant in ATP–binding cassette subfamily A member 7 (ABCA7), mid-exon frameshift (p.Leu1493ArgfsTer7), was identified in one SAD patient. ABCA7 is a membrane protein involved in lipid metabolism and phagocytosis^24^, and has been implicated in APP processing and Aβ clearance^25,26^. The frameshift found in ABCA7 occurs in the large extracellular loop between the first and second transmembrane helix in the second transmembrane domain. Several frameshifts identified in ABCA7 are associated with AD; however, the frameshift found in our sample set has not been implicated as a causal variant^27^. Our proteomic data nominates ABCA7 mid-exon frameshift p.Leu1493ArgfsTer7 as a candidate functional variant relevant to AD.

A striking difference in the case characteristics between the dominant-like and dominant-unlike subgroups of SAD in SMTG was age at death. For the dominant-like SAD subgroup that more closely resembles the ADAD SMTG proteome, the majority of cases (5/7) were early-onset cases with death before 70 years of age, with the other two dying at 87 and 88 years of age. The dominant-like SAD subgroup then was composed largely of earlier onset individuals with 5 of the 7 younger than 70 years at death^28^. We speculate that exclusion of comorbidities increased the likelihood for selection of people with more aggressive, early onset AD than what typically occurs in older adults who most commonly have a mixture of AD and common comorbidities conspiring to produce cognitive impairment^8^.

Prolific accumulation of both tau and Aβ is commonly indicative of worse cognitive impairment^29^. While these differences in SAD proteomes were mainly observed in the SMTG, that does not necessarily mean it is distinct to that brain region. Unfortunately, the lack of ADAD cases with available hippocampus material means we are unable to adequately examine the relationship between the ADAD and SAD proteomes. Based on the PCA clustering of the IPL, the Aβ and tau abundance differences, and peptide abundance differences found by testing with the sporadic cases classified into two categories, there are still differences in the SAD IPL proteome, however they are not as distinct as the SMTG.

Post-translational modification of tau is thought to affect its processing, with some modifications occurring more frequently in tau tangles in AD^30^. Modifications of tau have also been implicated in the rate of AD progression^31^. Multiple peptides mapping to residues 195-240 have decreased abundance in AD cases compared to the HCF cases. This is inverse of the peptides mapping to the microtubule-binding region, such as tau 354-369, which have the expected increased abundance in AD compared to the HCF cases. Since we are detecting and reporting the quantities for the unmodified tryptic peptides, the decrease we observe is likely due to a change in stoichiometry of the unmodified:modified peptides. If the majority of tau in the AD samples are hyperphosphorylated, then the stoichiometry would shift such that the unmodified peptide is relatively decreased. For all brain regions tryptic peptides significantly correlated with tau 354-369 have very similar trends to the peptides correlated with Aβ 17-28. The relationship of tau abundance in the SMTG and IPL with the subgroups of SAD supports the existing literature on the accumulation of both Aβ and tau together being more prominent in severe disease^32^.

Our findings highlight the importance of looking at different forms of AD to better understand how disease is related to aging. The striking differences within the SAD proteomes indicates the need for further study of the molecular differences between early onset SAD and late onset SAD with larger well characterized cohorts, like is being curated with the Longitudinal Early Onset Alzheimer’s Disease (LEADs) study^33^. Interestingly, for this current study, once the SAD cases were separated based on their SMTG proteome profiles rather than simply age, there were no longer many significant differences between the dominant-unlike SAD and RAD cases (no proteins or peptides differed significantly between them in SMTG). This is likely due in part to this current cohort being underpowered for uncovering molecular differences between these age-matched conditions^34^. However, the careful exclusion of comorbid neuropathology makes this a unique insight, and suggests that RAD is not only histopathologically but also proteomically similar to late onset, proteomically dominant-unlike SAD, and that unappreciated early-onset, proteomically dominant-like SAD cases could be contributing to the distinct molecular profiles seen in large cohort studies. Additional study into the late onset AD with genetic factors could also shed light on dissecting the role of age on disease neuropathology with differing etiologies^35^.

## METHODS

### Human brain sample collection and proteomic processing

Brain samples were collected under protocols approved by the Institutional Review Board (IRB) at University of Washington and collected through the Adult Changes in Thought (ACT) study, or at Washington University in St. Louis and collected through the Dominantly Inherited Alzheimer’s Network (DIAN). Cognitive status was diagnosed as dementia or not dementia by DSM-IVR criteria; and neuropathologic changes of AD were determined by consensus criteria; cases were excluded with any neuropathologic changes of Lewy body disease or pathologic TDP-43 inclusions in any region other than amygdala, any hemorrhage, any territorial or lacunar infarct, more than 2 cerebral microinfarcts, or hippocampal sclerosis. Tissue was cryopreserved from hippocampus by coronal section at the level of lateral geniculate nucleus, superior and medial temporal gyri (SMTG), inferior parietal lobule (IPL), and caudate nucleus at the level of the anterior commissure. For each of the four regions samples were divided into batches which included both inter-batch external quality controls and inter-brain region quality controls. Tissue was processed for mass spectrometry analysis as previously described^13^. In brief, two 25 μm frozen sections of brain tissue were homogenized and lysed using a Barocycler 2320EXT system (Pressure Biosciences Inc.), and protein digested with trypsin on S-trap columns (Protifi). Yeast enolase protein was added prior to digestion and heavy labeled Peptide Retention Time Calibrant (PRTC) mixture (Pierce) added post-digestion.

### Proteomic analysis

Specifics of the liquid chromatography and mass spectrometry acquisition are available in detail in Merrihew et al^13^. Briefly; one µg of each digested sample was separated by a Thermo Easy-nLC 1200 and analyzed by staggered window DIA on a Thermo Orbitrap Fusion Lumos Mass Spectrometer. Individual samples were acquired with 8*m/z* windows spanning 400-1000 m/z precursor mass range. Six gas-phase fractionated injections spanning the 400-100 m/z precursor mass range in 4m/z staggered windows were collected from pooled samples for use as an experiment specific peptide library as described previously^36^.

### Data processing

RAW files were demultiplexed and converted to mzML format using Proteowizard (version 3.0.20064). Peptides were detected with DIA-NN (version 2.6.1) using a Carafe fine-tuned, experiment-specific spectral library, and the DIA-NN results were imported into Skyline for transition selection and peak integration of each individual sample. Peptides were mapped to protein sequences from a Uniprot human canonical FASTA (January 2021), and peak integration was performed with Skyline. Additional manual peak integration was performed in Skyline for the peptides mapping to microtubule associated protein tau (MAPT; P10636). Peptide abundances were the sum of their transitions. For each brain region, data were normalized, batch corrected and rolled up to protein abundances with Skyline-PRISM (version 26.30.0): retention-time LOWESS normalization, ComBat batch correction at the peptide and protein levels, and Tukey median-polish protein rollup. Abundances were log2 transformed for all analyses.

### Statistical analysis

Peptide correlations with LVFFAEDVGSNK (APP residues 688-699; amyloid-β residues 17-28) for each region were calculated using a Spearman rank-based correlation, and significance determined as a Bonferroni adjusted p-value <0.05. Differential abundance testing was performed at the peptide and protein level with a moderated t-test (limma-trend empirical Bayes, with peptide count as the trend covariate at the protein level) for every pair of groups, using either the 4 clinical conditions or the 5 conditions generated from separating SAD into two subgroups. A feature was counted as significantly different when any pairwise contrast was significant. Significance was determined as a Benjamini-Hochberg adjusted p-value <0.05. Hierarchical clustering was performed with Ward linkage on z-score normalized peptide abundances. Principal component analysis was also performed on z-score normalized peptide abundances. Case characteristics were compared between SAD subgroups with Welch’s t-test (age, last MMSE and post-mortem interval), Fisher’s exact test (sex and APOE genotype) and the Mann-Whitney U test (B score, CERAD score, amyloid angiopathy and atherosclerosis). Enrichment analysis of gene ontology terms was performed using the Enrichr tool (GO Biological Process 2023)^37^, with biological process terms reported based on their adjusted p-values and low-overlap between terms (less than half of genes co-occurring with a similarly significant term).

### Exome sequencing and variant detection

Sporadic AD samples (specifically cases with SMTG tissue available in this cohort) were sequenced by Psomagen using a SureSelect V8-Post library kit and SureSelectXT Target Enrichment on an Illumina NovaSeq, multiplexed paired-end 151 base pair. The sequencing library was version C2 (Dec 2018). Sequence alignment and variant calling was performed for *APP, PSEN1, PSEN2, SORL1, ABCA17, TREM2,* and *ABI3* genes by the NW Genomics Center with bwa, GATK and the Seqr platform.

## DATA AVAILABILITY

Mass spectrometry data is available from Panorama Public at https://panoramaweb.org/ADBrainCleanDiagDIA.url the ProteomeXchange identifier PXD034525. Statistical analysis and code to recreate the figures is available on github at https://github.com/uw-maccosslab/manuscript-ad-subtype.

## ACKNOWLEDGEMENTS

Support for this research was provided by the National Institute of Aging (RF1 AG053959, U19 AG065156, F31 AG069420, R01 AG075802). The collection of tissue samples were supported by the UW Alzheimer’s Disease Research Center (P30 AG066509), the Adult Changes in Thought study U19 AG066567, the Nancy and Buster Alvord Endowment (C.D.K.), the Massachusetts Alzheimer Disease Research Center (P30 AG062421), The Dominantly Inherited Alzheimer Network (U19 AG032438, UF1AG032438), the German Center for Neurodegenerative Diseases (DZNE), Raul Carrea Institute for Neurological Research (FLENI), the Research and Development Grants for Dementia from Japan Agency for Medical Research and Development, AMED, and the Korea Health Technology R&D Project through the Korea Health Industry Development Institute (KHIDI). This manuscript has been reviewed by DIAN Study investigators for scientific content and consistency of data interpretation with previous DIAN Study publications. We acknowledge the altruism of the participants and their families and contributions of the research and support staff at each of the participating sites for their contributions to this study.

