## Supplemental Materials for "Peptide-centric proteomics reveals molecular subtypes of sporadic Alzheimer’s disease"

* Current Affiliation: ThermoFisher Scientific, Seattle, WA, United States
### Current Affiliation: Department of Pathology, University of California San Diego, CA, United States

**SUPPLEMENTAL MATERIAL**

**Supplemental Table 1.** Additional metadata statistics and testing for significance between the dominant-like SAD group 2 and the dominant-unlike SAD group 1.

| | **dominant-like SAD** n=7 | **dominant-unlike SAD** n=11 | **P-value** | **Significance test** |
|---|---|---|---|---|
| Sex [F (%)] | 2 (28.57%) | 5 (45.45%) | 0.64 | Fisher's Exact |
| Age [yrs (±SD)] | 69.57 (±12.49) | 89.18 (±5.96) | 0.0049 | Welch's T-test |
| Last MMSE | | | <0.0001 | Welch's T-test |
| Mean (SD) | 7.17 (±4.07) | 21.50 (±3.50) | | |
| Missing (%) | 1 (14.29%) | 1 (9.09%) | | |
| PMI [hrs (±SD)] | 4.25 (±0.87) | 4.72 (±1.44) | 0.40 | Welch's T-test |
| APOE genotype | | | 0.75 | Fisher's Exact |
| ε3/ε3 count (%) | 4 (57.14%) | 8 (72.73%) | | |
| ε3/ε4 count (%) | 2 (28.57%) | 3 (27.27%) | | |
| ε4/ε4 count (%) | 1 (14.29%) | 0 (0.00%) | | |
| B score [mean (±SD)] | 3.00 (±0.00) | 3.00 (±0.00) | n/a | Mann–Whitney U |
| CERAD score [mean (±SD)] | 3.00 (±0.00) | 2.73 (±0.47) | 0.16 | Mann–Whitney U |
| Amyloid angiopathy, occipital lobe [mean (±SD)] | 2.57 (±1.13) | 1.91 (±0.83) | 0.18 | Mann–Whitney U |
| Atherosclerosis [mean (±SD)] | 2.14 (±0.69) | 2.55 (±0.69) | 0.21 | Mann–Whitney U |

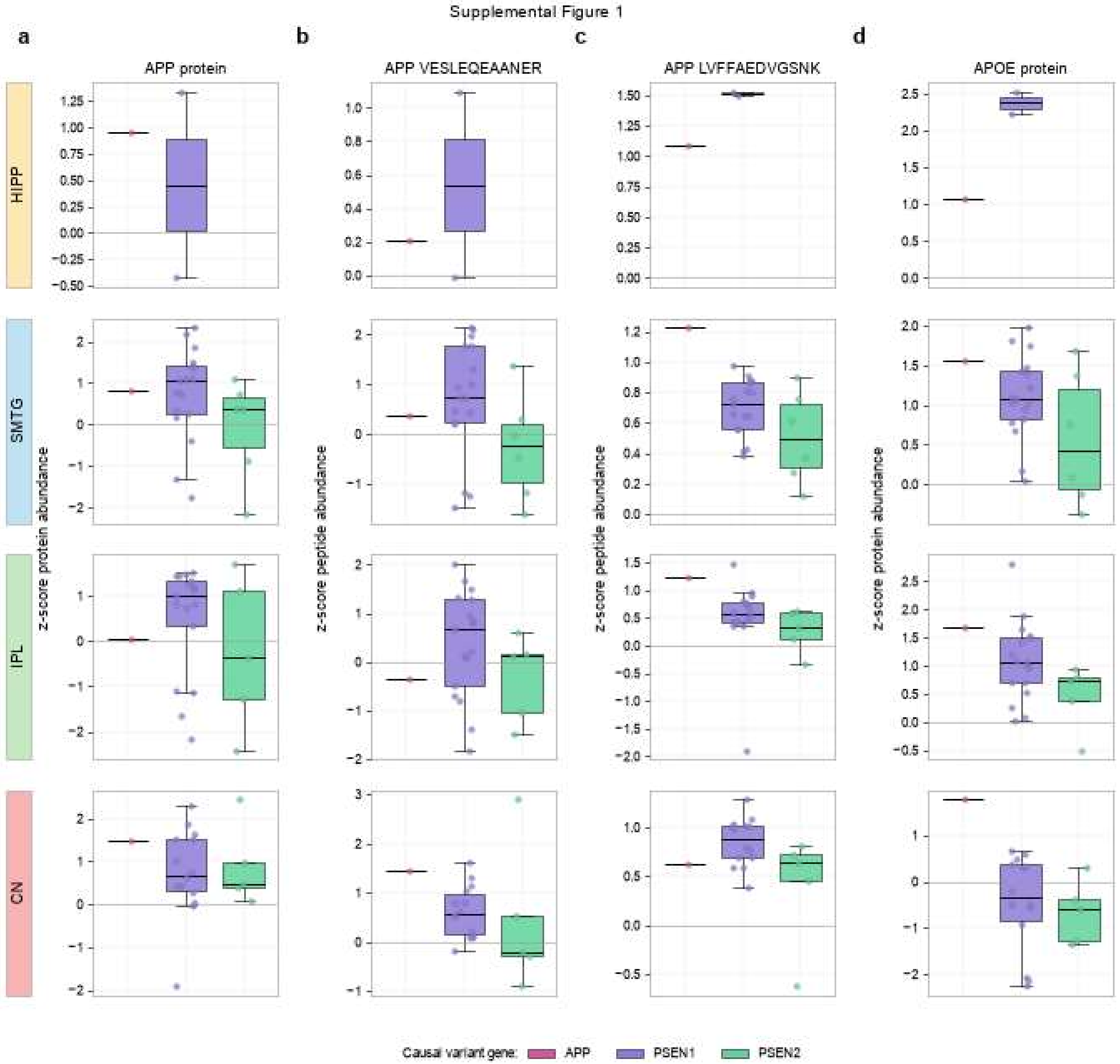


**Supplemental Figure 1. Protein and peptide abundance trends in autosomal dominant AD separated by causal variant gene.** A) Calculated APP protein abundance modestly increased in the cases with a causal variant in *PSEN1* compared to *PSEN2*. B) The same trend of an increase in the mean abundance is observed in APP peptide 439-450 in the E3 extracellular APP domain. C) Likewise, the same trend is observed in the Aβ 17-28 peptide, as well as observed in D) APOE protein abundance. (*APP* n=1, all regions; *PSEN1* Hipp n=2, SMTG n=11?, IPL n=13, caudate n=14; *PSEN2* SMTG n=6, IPL n=6, caudate n=5)

Supplemental Figure 2

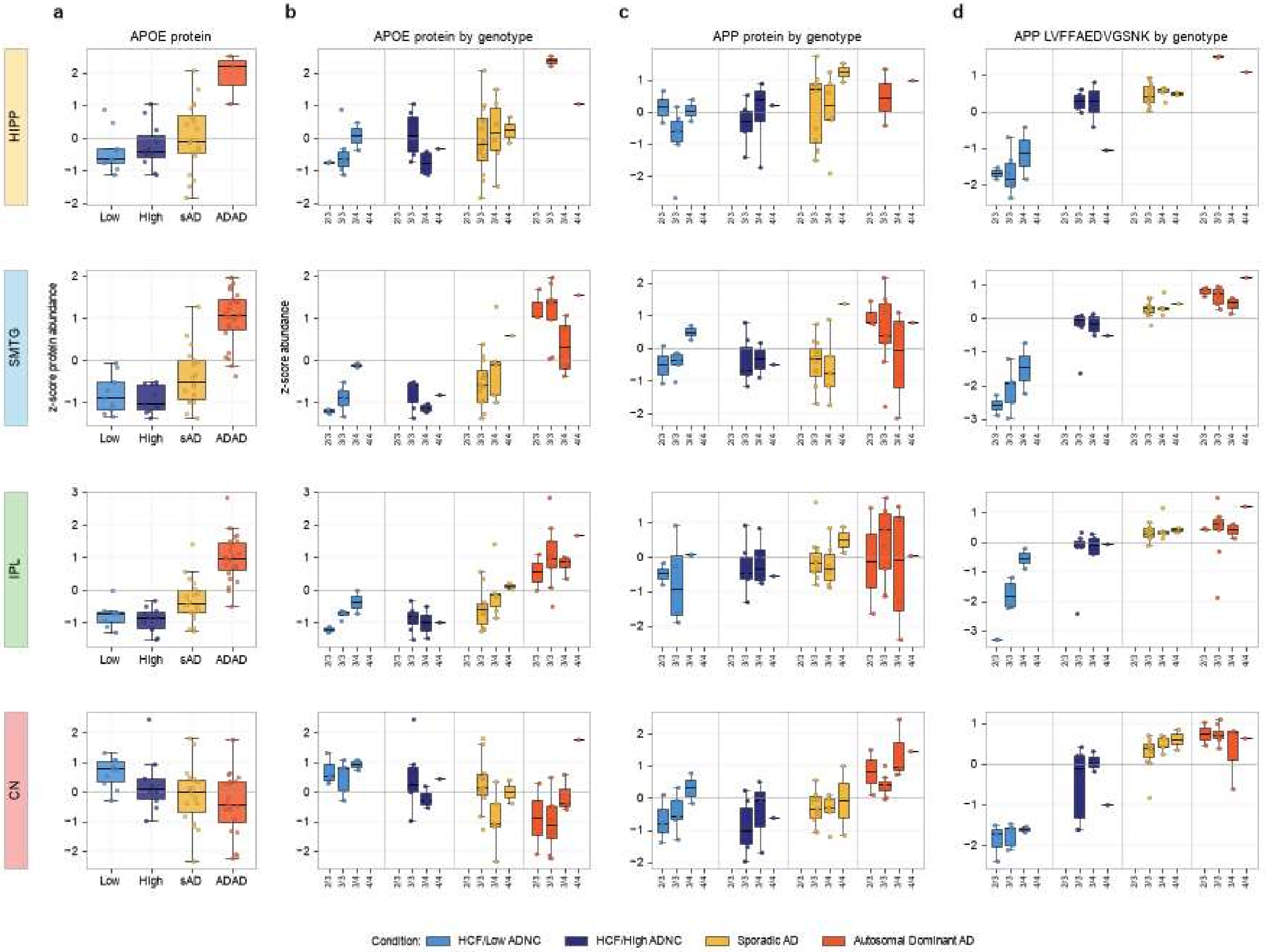


**Supplemental Figure 2. Protein and peptide abundance trends in APOE and APP by *APOE* genotype.** A) Normalized APOE protein abundance increased in autosomal dominant AD in all but the caudate nucleus region. B) The normalized APOE protein abundances for each condition separated by sample *APOE* genotype. C) Normalized APP protein abundance for each condition separated by sample *APOE* genotype. D) Normalized Aβ 17-28 tryptic peptide abundance for each condition separated by sample *APOE* genotype. Abundances are z-scored normalized across samples.

Supplemental Figure 3. All quantified tryptic peptides

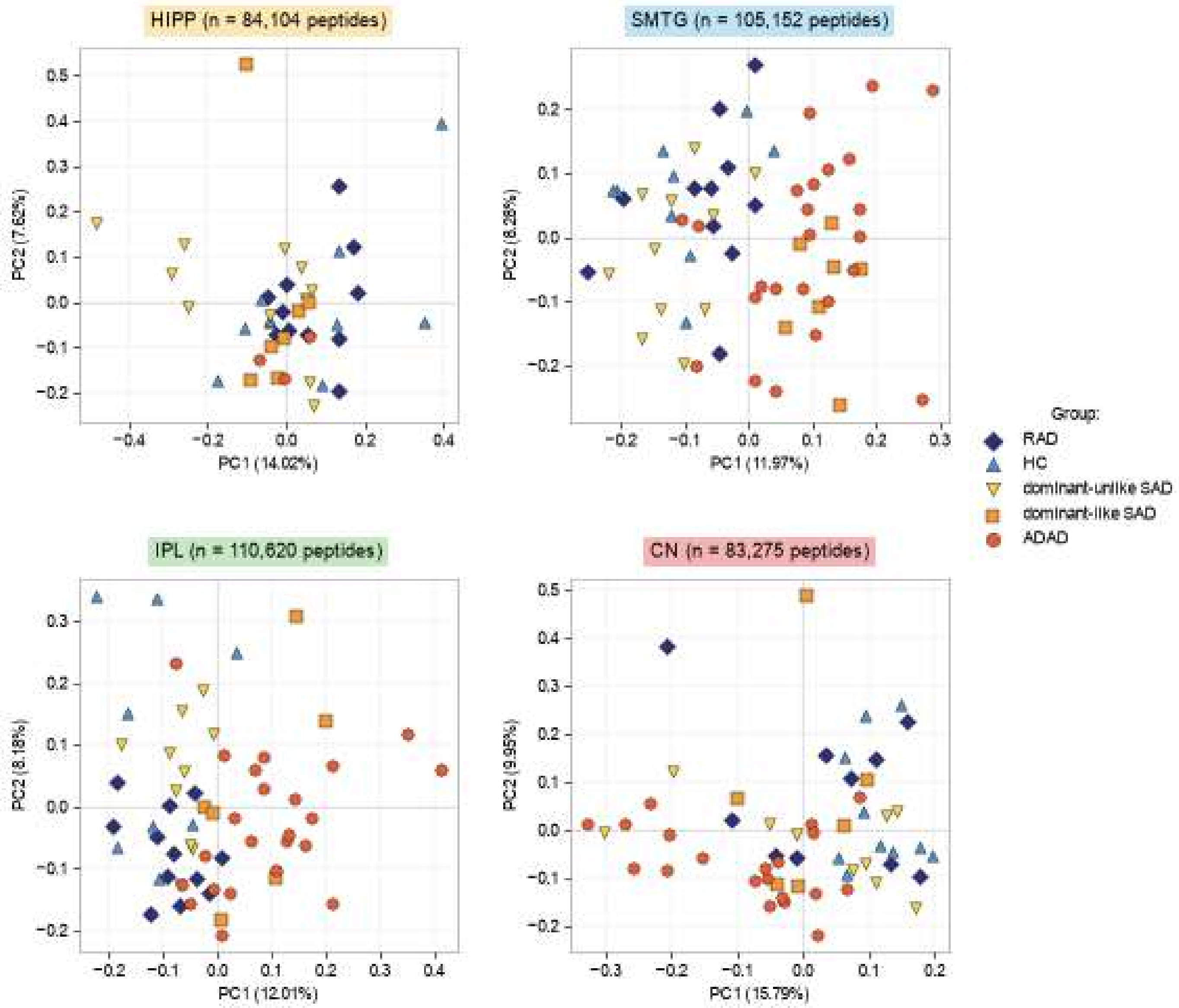


**Supplemental Figure 3. Principal component analysis.** All quantified tryptic peptides (hippocampus n=84,104, SMTG n=105,152, IPL n=110,620, caudate n=83,275).

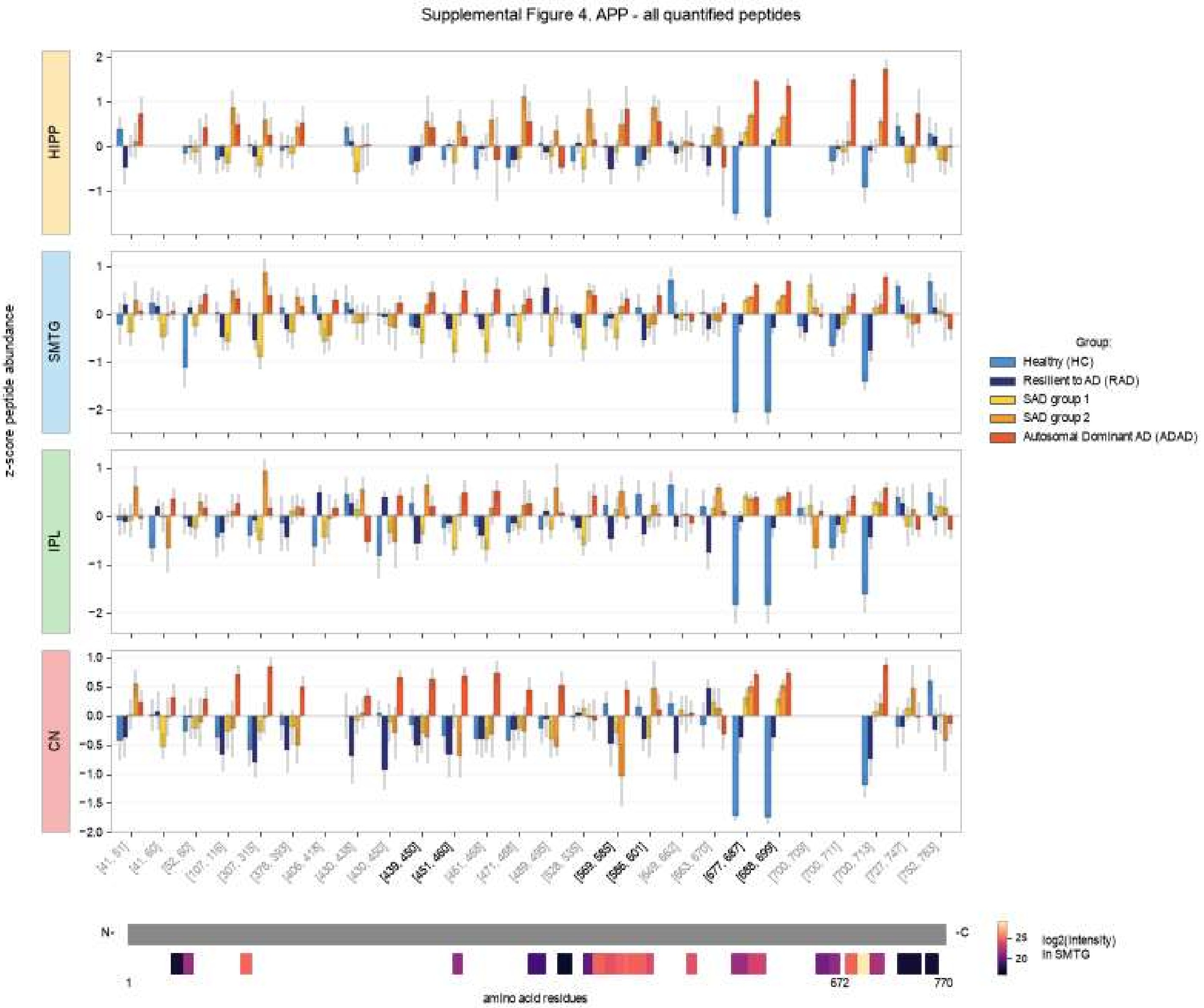


**Supplemental Figure 4. Additional APP tryptic peptide measurements**. All quantified tryptic peptides mapping to amyloid precursor protein (APP). Peptide z-score normalized abundance plotted by mean +/- se per condition from N- to C-terminus. Condition is indicated by color of the bar; with healthy control far left, then resilient to AD, SAD group 1, SAD group 2, and autosomal dominant AD. Peptides with grey residue numbers not included in main text figure.

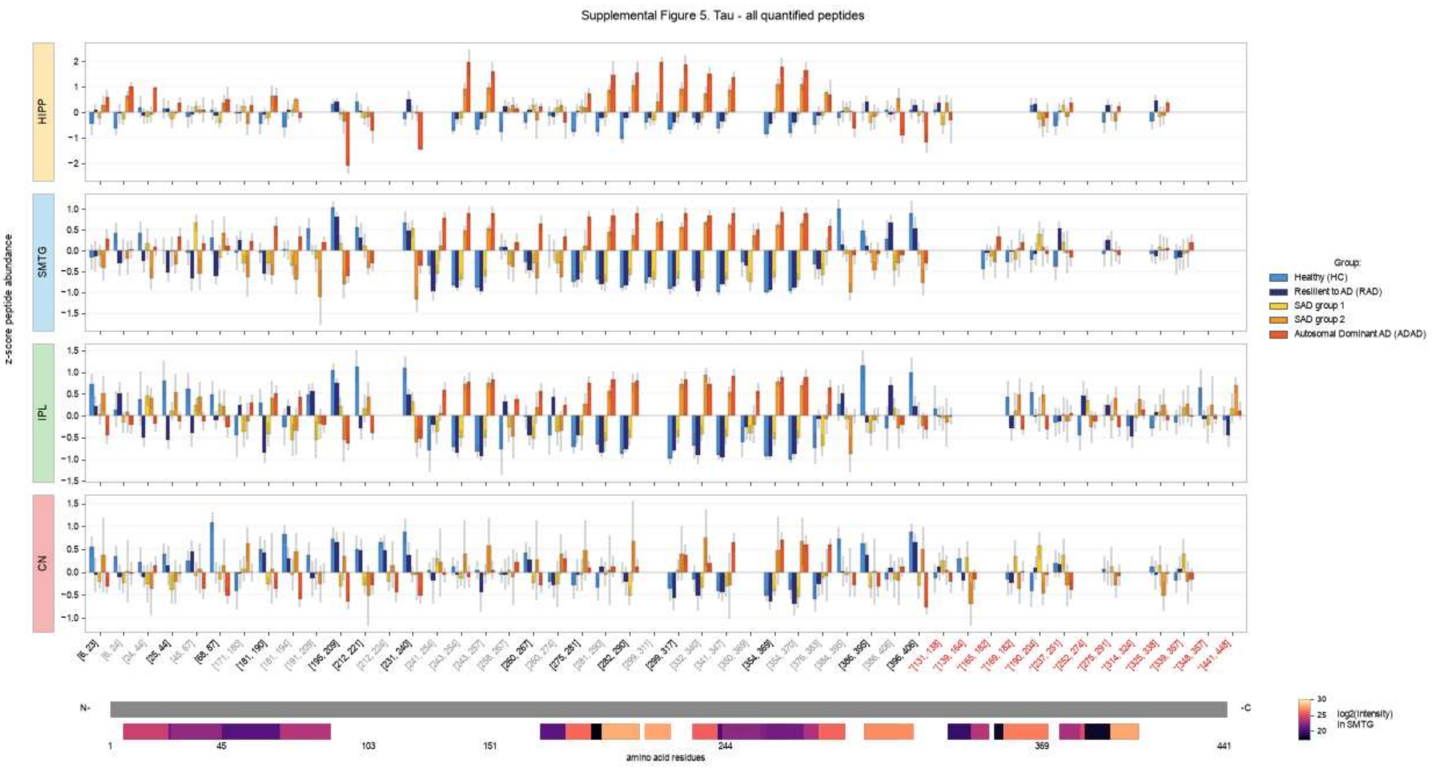


**Supplemental Figure 5. Additional TAU tryptic peptide measurements**. All quantified tryptic peptides mapping to microtubule associated protein tau (TAU). Peptide z-score normalized abundance plotted by mean +/- se per condition from N- to C-terminus. Peptides absent from the 2N4R isoform (red, marked *) are labeled with their residue numbers in the 758 aa canonical TAU sequence. Condition is indicated by color of the bar; with healthy control far left, then resilient to AD, SAD group 1, SAD group 2, and autosomal dominant AD. Peptides with grey residue numbers not included in main text figure.
